# Revisiting Color Efficient Coding through Material Perception

**DOI:** 10.1101/2025.03.22.644715

**Authors:** Sawayama Masataka

## Abstract

An essential objective of early visual processing is to handle the redundant inputs from natural environments efficiently. For instance, cone signals from natural environments are highly correlated across cone types, indicating channel redundancy. Early visual processing transforms the signals into color and luminance information, such as principle component analysis, and thus achieves efficient and decorrelated representations of natural scenes [1–3]. Building on these findings, previous research has investigated the effect of color on visual tasks such as object recognition using grayscale conversion, which separates luminance from color [4, 5]. However, recent work suggests that when focusing on object materials, color and luminance remain highly redundant due to complex optical properties [6, 7]. Although this finding indicates that there may be a more efficient decomposition of signals, the specific algorithms remain unknown. This study derives that a classic computer graphics algorithm, the median cut [8], offers a novel approach to enhance visual processing efficiency while capturing diagnostic features to separate material from object geometry information (Fig. 1a and 1b). Human behavioral experiments show that color reduction based on the algorithm disturbs material classification while preserving object recognition (Fig. 1c). These findings suggest that object geometric structures are available only from low-bit information. Finally, considering material information can be estimated from summary statistics of image sub-spaces, this study suggests an efficient decomposition of input color images (Fig. 1d).

## 1 Main Text

This perspective on efficient color processing is based on three key findings from material perception studies. The first finding is that color and luminance are highly redundant in certain materials, particularly in wet or translucent objects with subsurface scattering [6, 7, 9]. For example, there is a strong negative correlation between color saturation and luminance in some translucent objects, because light scattering in colored medium increases color saturation while reducing luminance. Additionally, simple grayscale conversion cannot reliably control material appearance, unlike object recognition. Specifically, for some objects, grayscale enhances translucency, while for others it diminishes [6, 7]. These observations suggest the potential for a more efficient decomposition than color and luminance separation.

The second finding is related to the importance of color entropy in material perception. This is demonstrated in the literature on wetness perception. In previous work, an image-based manipulation technique was developed to control the visual appearance of wetness [9]. The study found that this technique is effective only for images with high color hue entropy: for images with a limited number of color, it is difficult to control the appearance of wetness. Additionally, recent neurophysiological research suggests that some V4 neurons in monkeys selectively respond to heterogeneous hue variation within a receptive field, in addition to neurons selective for uniform hues [10]. These findings suggest that the visual system is sensitive to color varieties of a visual field and raise a possibility that the decomposition along the dimension of color varieties separates material from other object attributes.

The third finding is the presence of robust image features for shape perception that are independent of materials. As such a feature, Koenderink and van Doorn [11] derive that the geometric information of an object is inherited within a spatial map of image gradient directions (e.g., isophote map) of the image. One way to edit materials while keeping this feature is to manipulate the shape of the cumulative histogram of an image, which maintains the local intensity order [12]. For instance, when the cumulative histogram of a source image is matched to that of a target image, the material appearance of the source is transferred to the target. In addition to computational and behavioral findings, neurophysiology research has found some V4 neurons that selectively process shape information independent of material properties [13]. These findings suggest that image manipulation using cumulative histograms enables us to separately control materials from shapes.

These three key features are implicitly included in a classic color reduction method known as the median cut algorithm [8]. In summary: (1) the algorithm operates in a color space where color and luminance are mixed; (2) controls the number of color indices, (i.e., color variety), used to encode an image; and (3) manipulates the image using cumulative histograms of each input signal. The algorithm divides an input color space (Fig. 1b) according to a pre-defined number of color indices. It identifies the axis with the maximum variance from each RGB (or LMS) intensity histogram and selects the median of the cumulative histogram along that axis. Using this median value, the 3D color space is split along the selected axis, resulting in two subspaces. This process repeats until the number of subspaces equals the predefined number. Finally, the mean value within each subspace is applied to all points in that subspace (Fig. 1b, right). An image visualized using only four color indices can still convey the geometric structure of a scene because the local gradient direction remains relatively unchanged, though material appearance is lost.

**Fig. 1.**
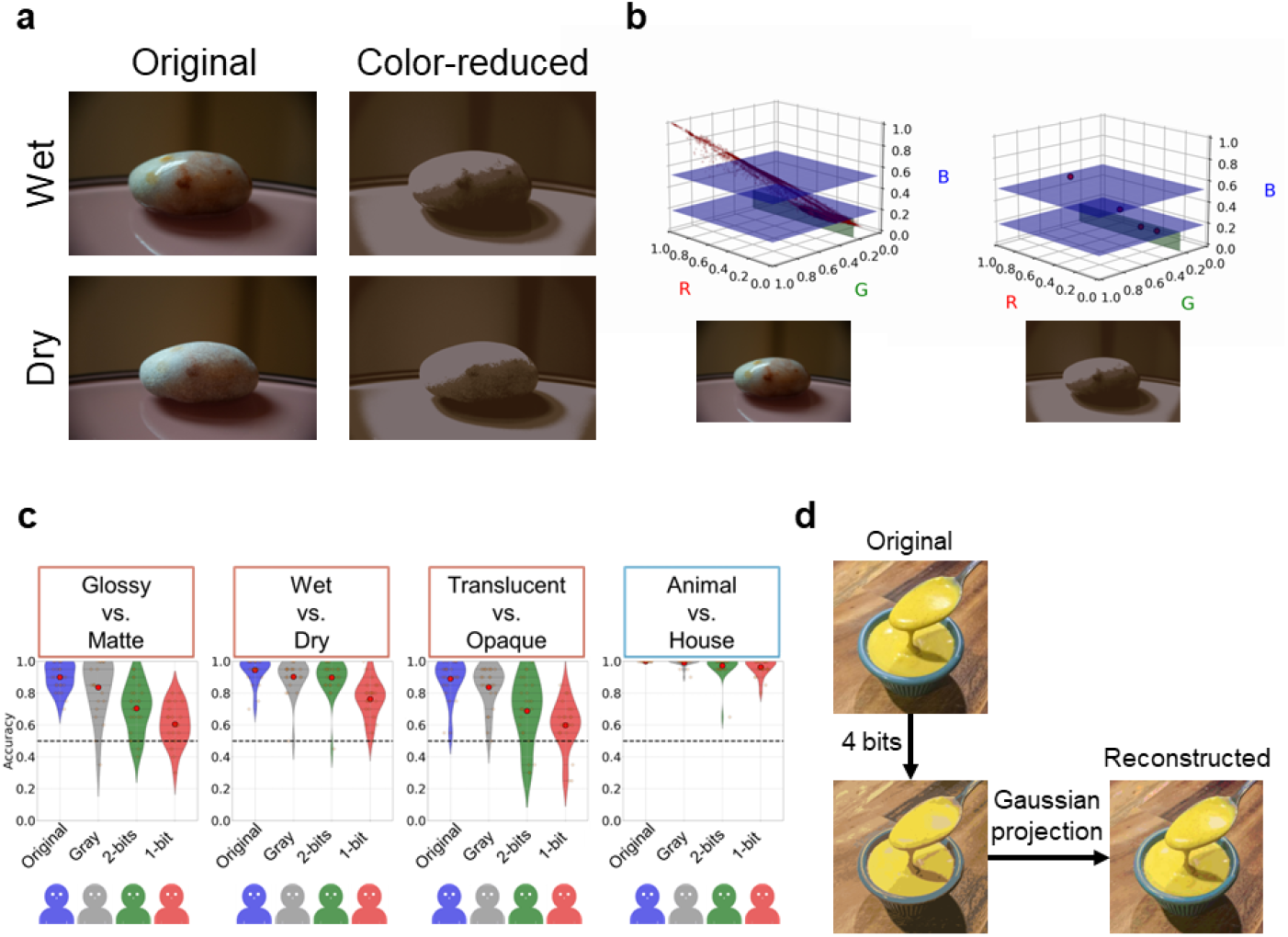
(a) Demonstration of canceling the material appearance difference (wet and dry) of the same stone image while maintaining its geometric structure using a color reduction algorithm called the median cut. The reduced color image consists of four color points in a RGB space, corresponding to 2-bits index color. (b) Schematic visualization of the median cut. The input color space is divided with a plane according to the median of the cumulative histogram of each axis. Each point in the left figure represents a pixel of the RGB color image. The divisions by the plane define the subspaces, and all points within each subspace are replaced with a single mean value. The right figure shows an example with 2-bits. According to the histogram of each subspace, the intensity order of the mean values is maintained consistent with the original distribution (Supplementary Fig. S1). (c) Results of human behavioral experiments. One hundred participants engaged in ensemble 2AFC tasks involving three material classifications and one object classification: Glossy vs. Matte, Wet vs. Dry, Translucent vs. Opaque for material classification, and Animal vs. House for object classification. Participants were divided into four groups based on the color condition to ensure that no participant saw the same original image. Violin plots in each panel illustrate the accuracy distribution for these image conditions, with the mean accuracy indicated by red dots. The horizontal dashed line marks the chance level accuracy. The results demonstrate that color reduction decreases material classification accuracy but does not affect object classification accuracy. (d) Color image decomposition by the median cut with Gaussian ellipsoid projection. While the median cut preserves the geometric structure of an image, the residual material information can be efficiently represented by the summary statistics of the Gaussian ellipsoid within each subspace, i.e., the mean and standard deviation.

Human material and object classifications were evaluated by behavioral experiments. Original images from the THINGS dataset [14] were used and converted to both grayscale and color-reduced formats (Fig. 1c). The results showed that color reduction, rather than grayscale conversion, disrupted material classification but did not affect object classification (Fig. 1c), consistent with that low-bit images retain object geometry information, although much of the material information is lost.

Finally, how can the residual material information be represented? One possibility is that it is represented by summary statistics of each 3D subspace(Fig. 1d). By assuming a Gaussian ellipsoid within each subspace of the median cut, material information is efficiently encoded with the standard deviation of each subspace.

This study proposes an optimal computational strategy, acknowledging that biological brains have evolved to efficiently process natural environments including materials. This work contributes to a novel hypothesis to uncover the still-unknown mechanisms of brain information processing. Additionally, it could impact the encoding process of most recent artificial intelligence models, which overlook the details of optical materials while training on natural object image datasets.

## 2 Supplemental Methods

### 2.1 Median cut algorithm

The median cut algorithm [8] can be applied to various color spaces, such as RGB or LMS. Figure S1 illustrates the algorithm using the RGB space of an image. The process begins by determining the number of color indices for encoding, which is set to four in this case. The algorithm then identifies the channel with the highest variance from the RGB intensity histograms; here, the B channel has the highest variance. For this channel, the cumulative histogram is calculated to locate the median of the distribution. Using this median value, the RGB space is divided at the B channel median, resulting in two 3D subspaces, lower and upper. This process repeats until the number of subspaces equals four. Notably, the mean distribution of each RGB channel in four 3D subspaces is sorted in the order of subspaces 1, 2, 3, and 4, respectively. This occurs because the cutting is based on the cumulative histogram, allowing the intensity order of the original histogram to be preserved. Although the cuts are applied to different channels depending on the subspace distributions, the input color space shows high redundancy in the cross-channel correlation of natural images. Therefore, the intensity order of each channel can be consistently maintained by the median cut. Consequently, in the spatial domain, the output image, visualized using only four color indices, still allows recognition of the scene structure since the local gradient direction of the image remains relatively unchanged (cf., [12]).

Figure S2 shows examples of varying the number of palette colors. In Fig. S2a, the wet and dry appearances become difficult to distinguish in images with a small number of colors. In Fig. S2b, as the number of colors decreases, the liquid appearance of the viscous mustard becomes ambiguous, and the glossy impression of the mustard and spoon fades.

### 2.2 Behavioral experiments

#### 2.2.1 Participants

One hundred naïve participants engaged in the experiments. The participants were recruited with Prolific (https://www.prolific.com/). All gave informed consent, which was approved by the Ethics Committee of the University of Tokyo.

#### 2.2.2 Stimuli

A major challenge in testing material appearance is the lack of large image datasets with diverse material attributes. For example, there are no publicly available big datasets of wet and dry images. To address this, we sampled material images from existing object image datasets using pre-trained vision and language models (Fig. S3a). Specifically, we leveraged the zero-shot image classification with pre-trained vision and language models [15]. In this method, a pair of pre-trained vision and text encoders are used. These encoders are trained to acquire shared representations of various text-image pairs, so we can input arbitrary text to find matching images without additional training, called zero-shot prediction.

Using a pair of pre-trained vision and text encoders, we input text prompts like “an object is X” to a text encoder. The word X is selected from “wet,” “dry,” “glossy,” “matte,” “translucent,” or “opaque” for material classification, and “animal” or “house” for object classification. Then, we input all the images of an object image dataset, THINGS database [14], to a corresponding image encoder to the text encoder. We computed the cosine similarity between the embeddings of the image and text encoders and extracted the most similar images to each text prompt. To mitigate estimation biases by a specific model, we employed this image extraction using six different vision-language models and averaged the cosine similarities of each image across the models. In the experiment, we used the top 85 images based on this extraction.

For each extracted image, we created four color conditions: original, grayscale, 2-bits by median cut, and 1-bit by median cut (Fig. S3b). For grayscale conversion, we assumed that each RGB image is recorded in linear sRGB color format. The median cut algorithm follows the above subsection.

#### 2.2.3 Procedure

One hundred participants were assigned to each condition with 25 in each, ensuring no participant saw the same original image (Fig. S3b). Each participant engaged in three material classification tasks and one object classification task in a random order. In the experiment, we adopted an ensemble judgment of material appearance to avoid category-specific responses (Figs. S3c and S3d). Participants might guess the material properties of an object if they recognize its category, e.g., the wet state of a dolphin. To address this, four image pairs of each class were randomly extracted from the 85 images and presented for 200ms in each trial. Participants fixated on the center probe of the display [16] and were asked to judge material or object classification with a two-alternative forced choice: glossy vs. matte, wet vs. dry, translucent vs. opaque for material classification, or animal vs. house for object classification. Each task was repeated 20 times, and all the images presented were different across trials.

Experiments were conducted online, with the code implemented using a JavaScript library (p5.js: https://p5js.org/) and a Python framework (Django: https://www.djangoproject.com/) [17].

### 2.3 Color image decomposition by the median cut with Gaussian ellipsoid projection

Based on the finding that the median cut method enables efficient color representation that can capture the geometry information of objects, this study also proposes a simple algorithm for representing the residual material information. The algorithm assumes a Gaussian sphere distribution for the subspaces segmented by the median cut in the input color space of an image. It then projects each point within these subspaces onto the surface of the Gaussian sphere.

In detail, the input RGB color space of an image is treated as a 3-dimensional space, where each pixel value is represented by a 3-dimensional vector **p** ∈ 𝒫 = [0, 1]^3^. The median cut method segments this space into multiple subspaces, denoted as 𝒮_*i*_, where *i* indexes the subspaces. Assuming a Gaussian ellipsoid for the distribution of pixels within each subspace, the projection vector **p**^*′*^ of a pixel **p** in a subspace 𝒮_*i*_ onto the Gaussian surface is given by:

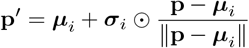

Here, ***µ***_*i*_ and ***σ***_*i*_ represent the mean and standard deviation vectors of the subspace distribution, respectively. Notably, because the pixel distribution within each sub-space is represented with summary statistics using the mean and standard deviation, reconstructing the image with a Gaussian ellipsoid provides an efficient representation. Figure 1d illustrates the reconstruction result using projections onto the surface of Gaussian ellipsoids, divided into 4 bits.

---

**Fig. S1.**
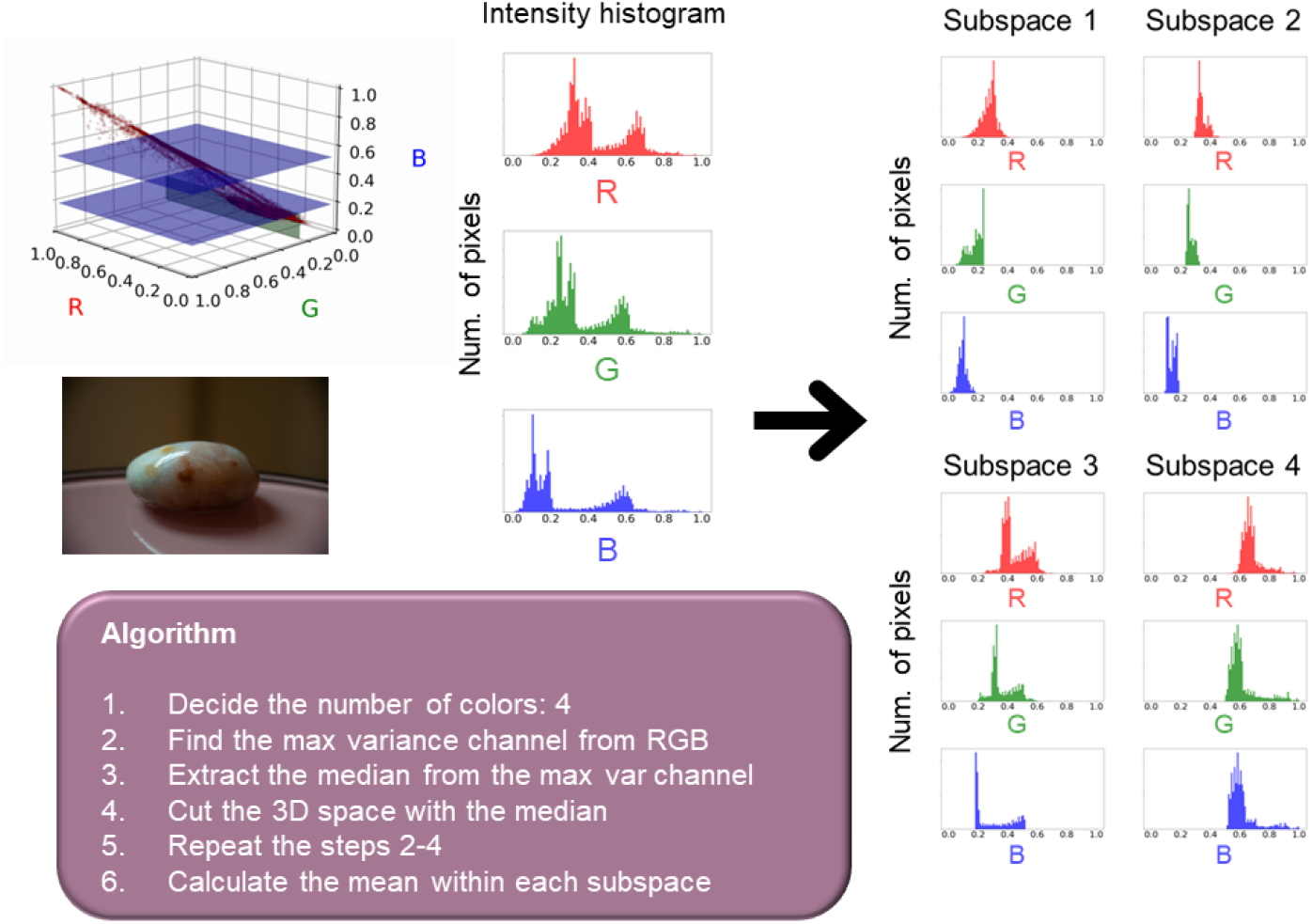
Overview of the median cut algorithm. The algorithm is applied to the input color space and aims to identify subspaces, as predefined by the user, to represent the original image with a limited number of color indices.

**Fig. S2.**
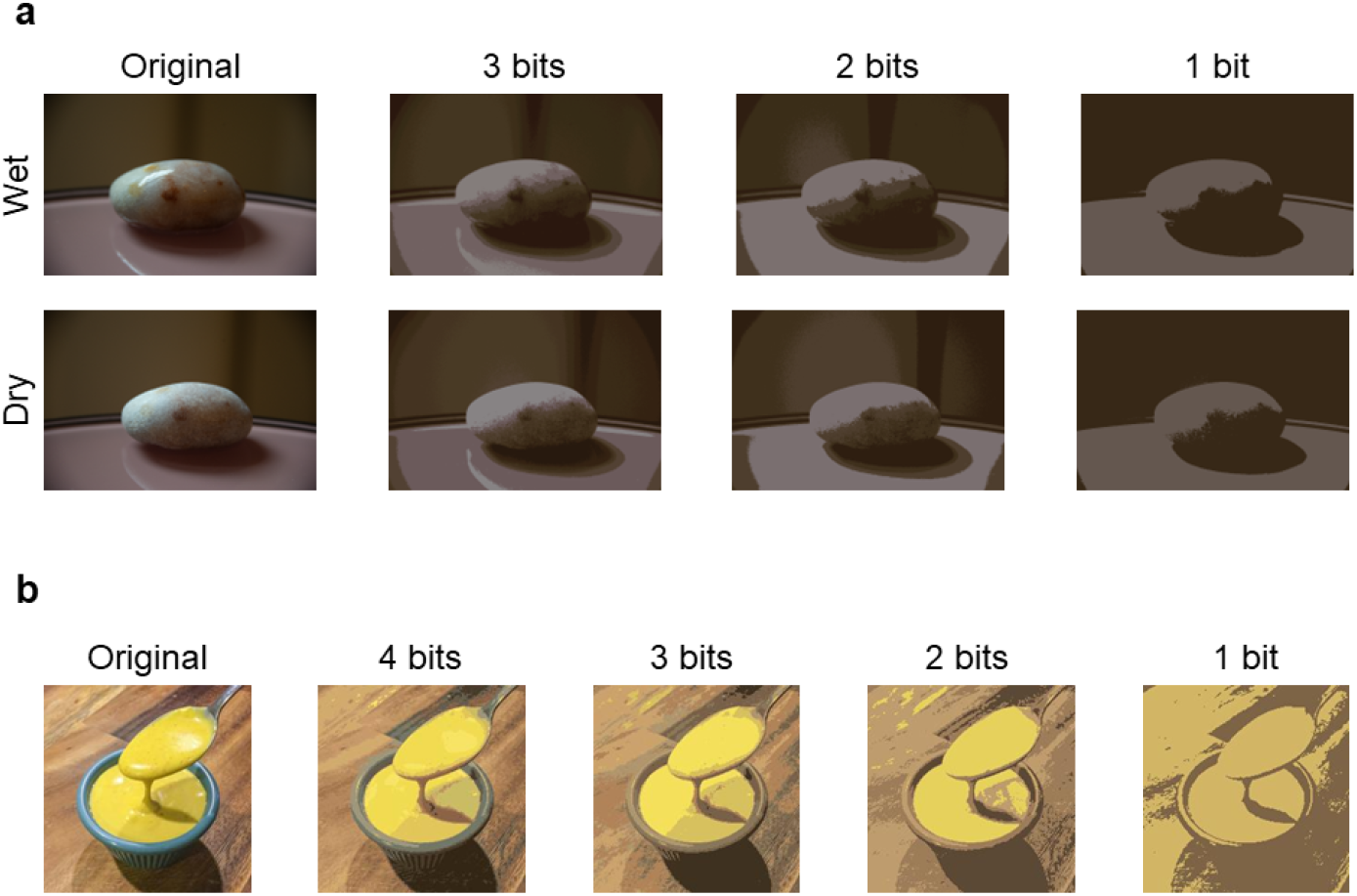
Examples of the median cut applied to material images. (a) Wet and dry stone images. The stone is the same object, photographed under the same conditions with different material states, i.e., wet or dry. (b) Viscous mustard image. The image is extracted from the THINGS dataset [14]. In both cases, as the number of colors decreases, the material appearance diminishes.

**Fig. S3.**
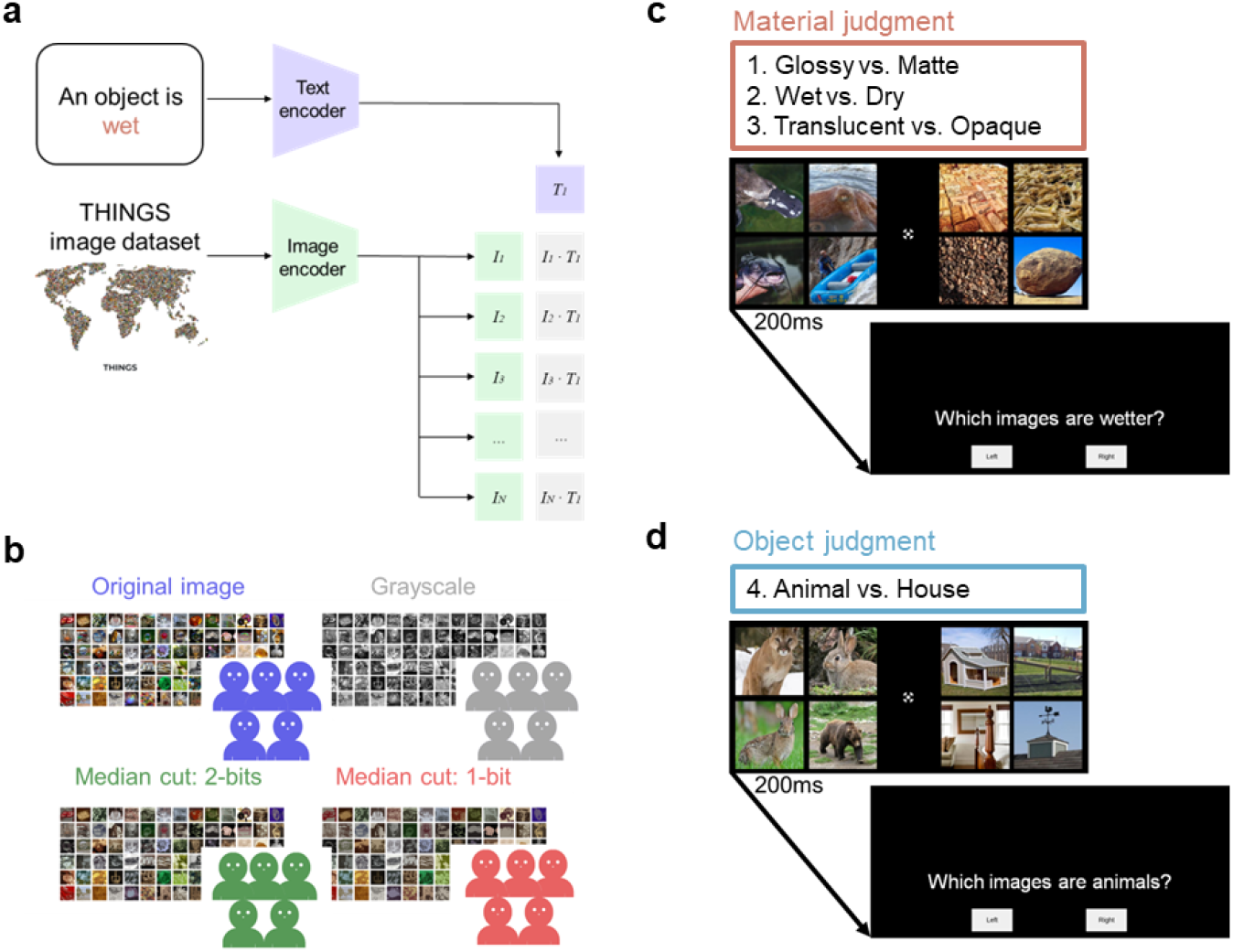
(a) Image sampling method for behavioral experiments. The illustration was created by editing the figures from [15] and [14]. Material images were extracted from the THINGS dataset using vision and language models. (b) The image color conditions used in the experiment: original, grayscale, 2-bits median cut, and 1-bit median cut. Different participants were assigned to each color condition in the experiment. (c, d) The time course of stimulus presentation. Each participant engaged in four tasks: three material judgment tasks and one object judgment task. In each task, a pair of four images was presented in the left and right visual fields. Participants were asked to identify the images with the target attributes.

